# Perceptual cycles travel across retinotopic space

**DOI:** 10.1101/2022.05.04.490030

**Authors:** Camille Fakche, Laura Dugué

## Abstract

Visual perception waxes and wanes periodically over time at low frequencies (theta, 4-7Hz; alpha, 8-13Hz) creating *perceptual cycles*. These perceptual cycles can be induced when stimulating the brain with a flickering visual stimulus at the theta or alpha frequency. Here, we took advantage of the well-known organization of the visual system into retinotopic maps (topographic correspondence between visual and cortical spaces) to assess the spatial organization of induced perceptual cycles. Specifically, we tested the hypothesis that they can propagate across the retinotopic space. A disk oscillating in luminance (inducer) at 4, 6, 8 or 10 Hz was presented in the periphery of the visual field to induce perceptual cycles at specific frequencies. Electroencephalographic (EEG) recordings verified that the brain responded at the corresponding inducer frequencies and their first harmonics. Perceptual cycles were assessed with a concurrent detection task –target stimuli were displayed at threshold contrast (50% detection) at random times during the inducer. Behavioral results confirmed that perceptual performance was modulated periodically by the inducer at each frequency. We additionally manipulated the distance between the target and the inducer (3 possible positions) and showed that the optimal phase, i.e., moment of highest target detection, shifted across target distance to the inducer, specifically when its flicker frequency was in the alpha range (8 and 10 Hz). These results demonstrate that induced alpha perceptual cycles travel across the retinotopic space in human at a propagation speed of 0.3-0.5 m/s, consistent with the speed of unmyelinated horizontal connections in the visual cortex.

## Introduction

Our perception phenomenologically appears continuous over time. Yet, our perceptual performance fluctuates periodically between favorable and less favorable moments (VanRullen and Koch, 2003, VanRullen, 2016). These *perceptual cycles* have been observed in human and non-human primates at low frequencies (theta, 4-7 Hz; alpha, 8-13 Hz) (Purves et al., 1996; VanRullen et al., 2005, 2006, 2007; Reddy et al., 2011; Landau and Fries, 2012; Fiebelkorn et al., 2013a; Song et al., 2014; Huang et al., 2015; Dugué et al., 2015, 2017; Senoussi et al., 2019; Michel et al., 2022; for review, see Kienitz et al., 2021; Re et al., 2023), and can be induced by flickering visual stimuli (Stefanics et al., 2010; Mathewson et al., 2012; Henry and Obleser, 2012; De Graaf et al., 2013; Henry et al., 2014; Spaak et al., 2014; Ten Oever and Sack, 2015; Kizuk and Mathewson, 2017). Electrophysiological recordings showed that perceptual cycles correlate with brain oscillations, and specifically their phase, at the same frequencies (Valera et al., 1981; Busch et al., 2009; Mathewson et al., 2009; Busch and VanRullen, 2010; Dugué et al., 2011; Fiebelkorn et al., 2013b; Hanslmayr et al., 2013; Samaha et al., 2017; Fakche et al., 2022). Yet, this phase effect accounts for less than 20% of the trial-by-trial variability in perceptual performance (Busch et al., 2009; Busch and VanRullen, 2010; Dugué et al., 2011, 2015; Baumgarten et al., 2015; Samaha et al., 2017; Fakche et al., 2022). Here, we build upon the well-known organization of the visual system and propose to fill this gap by assessing the spatial organization of induced perceptual cycles.

Visual input is topographically decomposed in the visual cortex, i.e., there is a topographic correspondence between the visual space and the cortical space called retinotopy (for review, Wandell and Winawer, 2011). Neural populations in the retinotopic cortex are connected by intra-cortical networks of long-range unmyelinated horizontal axons of up to several millimeters (for review, Chavane et al., 2022). Interestingly, invasive recordings in animals showed that brain activity originating from a specific location in the retinotopic cortex can propagate across the map (e.g., Bringuier et al., 1999; Jancke et al., 2004; Benucci et al., 2007; Muller et al., 2014; Zhang et al., 2012; Yang et al., 2015; Zanos et al., 2015; Townsend et al., 2015; Chemla et al., 2019; Davis et al., 2020). Here, we test the hypothesis that induced perceptual cycles can propagate across the retinotopic visual space presumably due to horizontal connections in the retinotopic visual cortex.

Sokoliuk and VanRullen (2016) showed in a psychophysics experiment that induced 5 and 10 Hz perceptual cycles assessed by detection of a low-contrast target stimulus propagated across the retinotopic visual space. Specifically, a brief target at threshold contrast was presented at three possible distances from a continuously oscillating disk in the periphery of the visual field (inducer). They showed that the optimal behavioral phase (the moment of the oscillating disk leading to the highest performance in target detection) shifted as a function of distance between target and inducer. Here, we used the same psychophysics approach. We additionally recorded EEG to identify the shape of the induced-brain activity, and eye-tracking to ensure successful fixation of the center of the screen, critical for stable retinotopic representation. Other critical changes were made to the design, including the size of the targets, which was adjusted to cortical magnification, so that each stimulus activated the same number of neurons in the retinotopic cortex. Additionally, we parametrized the frequencies by testing four different frequencies in the theta/alpha range (4, 6, 8, and 10 Hz), which was done in each participant (n=17) to ensure robust statistical analyses (in Sokoliuk and VanRullen, n=5 for 5 Hz, n=7 for 10 Hz and n=15 for a 10-Hz replication set). Finally, we added a control experiment to ensure that the results were not due to low-level luminance masking (in Sokoliuk and VanRullen, the control that was used was designed in a way that could have interacted with the attention of the participant).

Our study assessed the propagation of induced perceptual cycles across the retinotopic space. Contrary to the earlier study (Sokoliuk and VanRullen, 2016), our results demonstrate that alpha-induced perceptual cycles, and not theta, traveled across the retinotopic space in human observers at a propagation speed ranging from 0.3 to 0.5 m/s, consistent with the conduction delay of long-range horizontal connections in the visual cortex (Shao and Burkhalter, 1996; Nowak and Bullier, 1996; Girard et al., 2001).

## Materials and Methods

### Participants

18 participants (9 females, 16 right-handed, mean ± standard deviation (std) age: 26.2 ± 4.1 years) were recruited for the main experiment. One participant was excluded from the analysis because the number of false alarms exceeded the number of targets presented. 15 participants (7 females, 13 right-handed, mean ± sd age: 25.5 ± 3 years) were recruited for the additional control experiment 1 (see Flashing Disk section below), and 15 participants (9 females, mean ± sd age: 26.1 ± 4.3 years) for control experiment 2 (see Broadband Oscillating Disk). The number of participants was decided based on previous studies using flickering visual stimuli to induce perceptual cycles (mean ± sd number of participants from a sample of representative studies: 15.5 ± 9.3; Stefanics et al., 2010 (n=11; n=13); Mathewson et al., 2012 (n=13); Henry and Obleser, 2012 (n=12); Graaf et al., 2013 (n=21, n=18); Henry et al., 2014 (n=17); Spaak et al., 2014 (n=19); ten Oever and Sack, 2015 (n=20); Sokoliuk and VanRullen, 2016 (n=7, n=15, n=5, n=4); Kizuk and Mathewson, 2017 (n=42)). All participants were free from medication affecting the central nervous system, reported no history of psychiatric or neurological disorders, gave their written informed consent and were compensated for their participation. The study was approved by the local French ethics committee Ouest IV (IRB #20.04.16.71057) and followed the Declaration of Helsinki.

### Stimuli

Stimuli were designed with PsychToolbox 3.0.12, running in Matlab R2014b 64-bit (The MathWorks, Natick, MA), and displayed with a ProPixx Projector (Vpixx Technologies, Saint-Bruno, QC, Canada), on a 139 x 77.5 cm projection screen (960 x 540 pixels; 480 Hz refresh rate), at 122 cm distance. Three different stimuli were generated: a fixation cross, a peripheral disk, and three small dots (targets). The arms of the fixation cross measured 0.15° of length and 0.03° of width. The size and position of the targets were computed according to cortical magnification (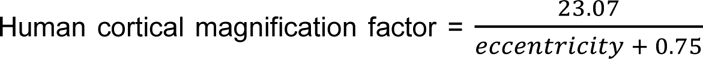; Horton and Hoyt, 1991) so that each target (diameter) represented a 0.8 mm portion of cortex spaced by 0.8 mm edge-to-edge (radius: 0.09°, 0.1° and 0.12°; eccentricity: 4.1°, 4.5° and 4.9°). The peripheral disk (inducer) was displayed on the lower right corner at 7.5° of eccentricity (from the center of the disk; radius: 1.75°). The background color was black, and the fixation cross at 50% of contrast relative to the background. Target contrasts were titrated for each participant, each position, and each frequency, using a staircase procedure (see below). The peripheral disk was oscillating sinusoidally in luminance, from black to white (i.e., from 0% to 100% of contrast; one period: black-to-white-to-black), at frequencies in the theta and alpha range: 4.07, 6.13, 8.27 and 10.43 Hz. For clarity, we rounded these values to 4, 6, 8, and 10 Hz.

### Eye tracker

Participant’s head was maintained with a headrest and chinrest. An infrared video-camera system (EyeLink 1000 plus 5.08, SR Research, Ottawa, Canada) was used to ensure that participants maintained their gaze on the fixation cross throughout the block. The block only started when participants were successfully maintaining fixation. When a gaze deviation of >2° from the center of the screen was detected during the presentation of a target (-150 ms to +100 ms around target onset) or a blink, we considered that the participant broke fixation and the trial was removed from the analysis (mean ± std: 129.1 ± 120.9 trials on average across participants, leading to a total number of trials ± std of 5015.2 ± 199.9 trials per participant, for the main experiment). Supernumerary blocks were added at the end of each run to compensate for rejected trials. Participants received feedback at the end of each block to remind them to minimize blinking and to maintain fixation.

### EEG

EEG was recorded using a 64-channels actiChamp system (Brain Products GmbH). The ground was placed at the Fpz position, and the right mastoid was used as reference (DC recording; 1000 Hz sampling rate).

### Experimental Design

Participants performed five sessions: four psychophysics sessions (one session for each induced frequency; frequency order randomized across participants), and one EEG session. The psychophysics sessions were composed of the staircases and two runs of 50 blocks each. The EEG session contained four runs of 24 blocks each (one run for each induced frequency).

For both the staircases and the main task, each block lasted 30 seconds during which the peripheral disk continuously oscillated in luminance. 6 to 18 targets were presented at random times (excluding the first and the last seconds, and with at least one second interval between targets) during three frames (6.3 ms) according to a decreasing probability distribution (∼10 targets per block on average). The number of targets for the three positions was randomized across blocks (and pseudorandomized across runs). Participants were instructed to press the space bar when they detected a target (in a 1-second time window after which their response was not considered; target presentation and response window composed a trial).

A one-up/one-down staircase with decreasing steps was performed separately for each target position to titrate the contrast of the target to reach about 50% detection. Each staircase was composed of 7 blocks, as described above, except that the targets appeared always at the same position. During the main task, target contrasts were adjusted every 15 blocks by multiplying the target contrasts to a factor y varying between 0.5 and 1.5 based on the detection performance for each target. y is calculated as follows

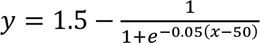

with x the detection performance. This procedure maintained detection performance at the same averaged level across the entire session. Target contrasts averaged across participants were mean ± standard error of the mean (sem): 3.2 ± 0.45 %, 2.2 ± 0.27 %, 2.88 ± 0.38 %, for position 1, 2, and 3, respectively, for the 4 Hz-induced frequency; 3.35 ± 0.55 %, 2.2 ± 0.29 %, 2.68 ± 0.27 %, for position 1, 2, and 3, respectively, for 6 Hz; 3.16 ± 0.31 %, 2.1 ± 0.19 %, 2.55 ± 0.18 %, for position 1, 2, and 3, respectively, for 8 Hz; and 3.03 ± 0.31 %, 2.02 ± 0.17 %, 2.45 ± 0.16 %, for position 1, 2, and 3, respectively, for 10 Hz.

### EEG Analysis

Analyses were performed with EEGLab 13.6.5 (Delorme and Makeig, 2004; Swartz Center for Computational Neuroscience, UC San Diego, California) running in Matlab.

#### Preprocessing

EEG data and channel location were imported into EEGLab. EEG data were re-referenced to the average of the right and left mastoids. A high-pass filter at 0.1 Hz and a notch filter between 48 and 52 Hz were applied, to respectively remove slow drifts and 50 Hz electric noise. The signal was further down sampled to 512 Hz. Visual inspection allowed to identify electrodes with low signal-to-noise ratio, which were then interpolated (spherical interpolation). Independent component analysis (ICA) was performed to remove blink components, after a visual inspection of their time courses and topographies. Data were epoched from trial onset (0s) to the end of the block (+30s).

#### Fast Fourier Transform

FFT were computed (Matlab function: fft) on 30s-epoched EEG data, and the amplitude was extracted for each electrode, epoch, and frequency. The signal-to-noise ratio (SNR) was computed on the amplitude extracted from the FFT, for each electrode, as the ratio of the amplitude at each frequency to the average of the 20 surroundings frequencies (10 frequencies on either side, excluding the immediately adjacent frequencies; note that the SNR could mathematically not be estimated at the edges of the spectra), as described in Rossion et al. (2012) and Liu-Shang et al. (2014). SNR averaged across participants, epochs and electrodes were plotted to ensure that the oscillating disk successfully induced brain activity at the corresponding frequency. Note that an SNR > 1 indicates that there is a difference between the amplitude of the EEG signal at a given frequency and its neighboring ones. Topographies were plotted at the peak of the induced frequency and its first harmonic.

#### Event-Related Potentials (ERPs)

Previously preprocessed EEG timeseries were further bandpass filtered between 1.5 Hz and 30 Hz, and baseline corrected from -400 to 0 ms from block onset. Epochs of one second were defined (time-locked to full-black disk), excluding the first second of each 30 seconds-block to avoid the transient EEG response due to stimulus onset. Participants’ ERPs were computed as the averaged of all resulting epochs. ERPs averaged across participants for electrode Oz were plotted with the standard error of the mean. Electrode Oz was selected because it is the electrode with the highest amplitude across the four induced frequencies. The ERP plots allowed us to identify that the induced brain activity was composed of two sinusoidal brain signals, one at the induced frequency and the second at its first harmonic. Therefore, a complex sine function was fitted to the behavioral data.

#### FFT on ERPs

FFT was computed (1500 points zero padding) on the 1s-Oz ERP for each participant. The amplitude was extracted, and the SNR was computed as previously described.

### Behavioral analysis

Behavioral analyses were performed with Matlab R2014b (The MathWorks, Natick, MA). The following dependent variables were computed: hit rates as main dependent variable, i.e., percentage of correct responses, and median reaction times as secondary dependent variables, for each target position and frequency. Hit rates averaged across frequencies were: 47.65 ± 0.85 % (mean ± sem), 46.12 ± 0.81 %, and 45.02 ± 0.79 % for position 1, 2, and 3, respectively. Median reaction times averaged across frequencies were 512 ± 7 ms (mean ± sem), 512 ± 7 ms, and 512 ± 8 ms for position 1, 2, and 3, respectively. A two-way repeated-measures ANOVA was performed for each dependent variable to assess the effect of frequency and target position. For hit rates, there was a main effect of target position (F(2, 32) = 11.46, SS = 236.74, p-value < 0.01, eta^2^ = 41.75) but no effect of frequency (F(3, 48) = 1.58, SS = 76.31, p-value = 0.2, eta^2^ = 8.99) nor of their interaction (F(6, 96) = 0.36, SS = 12.66, p-value = 0.9). For median reaction times, there was no main effect of frequency (F(3, 48) = 2.08, SS = 4093.43, p-value = 0.11, eta^2^ = 11.55), nor main effect of target position (F(2, 32) = 0.04, SS = 12.38, p-value = 0.96, eta^2^ = 0.25), nor of their interaction (F(6, 96) = 1.02, SS = 622.66, p-value = 0.41). The absence of significant interaction between frequency and target positions in any of these two tests confirm successful contrast manipulation, i.e., changes in detection performance across target positions are not different across frequencies (in the absence of such interaction, main effects will not be further interpreted; importantly, the optimal phase can be properly estimated as performance did not reach ceiling or floor for any of the conditions, which was the purpose of the staircase procedure). As argued in Kienitz et al. (2022) and VanRullen and Dubois (2011), reaction time fluctuations do not unambiguously demonstrate rhythms in cognitive processes as they can also be the by-product of the external stimulation and are sensitive to changes in decision criteria (Reed, 1973; Wickelgren, 1977; Carrasco and McElree, 2001; Dugué et al., 2018, 2020). The behavioral phase analyses thus focused on hit rates as the main dependent variable.

#### Phase effect on detection performance

Each target was assigned to one of 7 phase bins of the periodic stimulation depending on the delay at which they appeared during the block. There were 124 blocks with on average 10 targets per block for each frequency, i.e., ∼1240 trials per frequency in total, resulting in ∼413 trials per position and ∼60 trials per phase bin. Hit and false alarm rates were computed for each target position, frequency, and phase bin. For each block and target position, false alarms (participants reported perceiving a target while no target had been presented; participants were instructed to respond in the 1s-window following the target, after which the response was considered a false alarm) were binned as a function of the phase of the oscillating disk, i.e., the phase of the false alarms depended on the phase bin of the participant’s response. To allow for a fair comparison between hit and false alarms, only the false alarms that were in the same phase bins as the targets (but in a different 100-ms period) were considered for further analysis (e.g., if 2 targets were presented at position 1 within the given block, and binned in bins number 2 and 6, only false alarms that were binned in bins number 2 and 6 were considered for further analysis). To allocate a false alarm to one of the three target positions (a false alarm is, by definition, at none of the position), a bootstrap procedure was performed (100 repetitions).

Given the ERP results (see **Figure 2**) showing a complex neural response composed of the induced frequency and its first harmonic, a complex sine function was fitted to behavioral data (Eq1) with the following free parameters: *x*[1] and *x*[2] the amplitude and the phase offset of the induced frequency, respectively, *x*[3] and *x*[4] the amplitude and the phase offset of the first harmonic, respectively, and *x*[5] the baseline level. Note that, since we did not have a priori hypothesis regarding the shape of the induced behavioral response, we selected the shape of the closest proxy, i.e., the evoked response. To find the parameters that best fit the data, we used a Least Squares cost function, i.e., the cost function computed the total squared errors, i.e., the sum of squared errors, between the fit and the data, thus giving a measure of how well the data are fitted. The best fit is the one whose parameters minimize the total error (Watt et al., 2020).

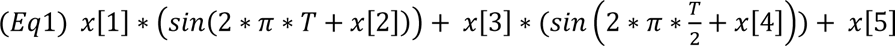

We then performed a Monte Carlo procedure on the fitted amplitude (see **Statistical Analyses** section) to evaluate whether the fitted data was modulated periodically as a function of the phase. Note that false alarms were low in each phase bins (< 0.5%) and not significantly modulated periodically after Bonferroni correction across frequencies and positions (p-value threshold < 0.004, corresponding to a Z-score threshold of 2.64, and a Cohen’s d threshold of 1.66). Consequently, in the next steps of the analysis, only hit rates were considered.

A group level analysis was also performed. Empirical and surrogate data were averaged across participants, and the same Monte Carlo procedure was performed.

#### Variance explained by the spatial organization of targets

We hypothesized that taking into consideration the spatial organization of induced perceptual cycles should explain more variance in the behavioral phase effect, i.e., the difference of performance between the optimal and the non-optimal behavioral phase should be larger when considering the spatial location of the target rather than disregarding where the target appears relative to the luminance disk. To test this hypothesis, we compared the amplitude modulation (difference in hit rate between the optimal and the non-optimal behavioral phase) averaged across the three target positions with the amplitude modulation computed across targets independently of their position, for each frequency separately. Specifically, hit rates were computed for each participant, frequency and phase bin, regardless of target position. We used a subsampling procedure (1,000 repetitions) to equalize the number of trials (pooled across all three target positions) to the number of trials for each position. Two-tailed t-tests were used to assess whether the amplitude modulation averaged across the three target positions was significantly different from the amplitude modulation computed across targets independently of their position, for each frequency.

#### Behavioral optimal phase shift between target positions

The optimal phase, i.e., the position on the fitted curve at maximal performance, was extracted for each target position and frequency, on data averaged across participants and for each participant individually. We asked whether the optimal phase shift as a function of target positions was different between the four induced frequency conditions, i.e., whether there is a significant interaction between frequency and target position.

The optimal phases were converted in radians. A one-cycle sine wave was generated with the phase offset of the oscillating disk and phase values were extracted in radians along the entire cycle. The optimal phase extracted from data fitting was then calculated in radian based on this one-cycle sine wave fit participant-by-participant. We then obtained rose plots of all individual optimal phases with the function circ_plot from the Circular Statistics Toolbox (see **Figure 4**; P. Berens, CircStat: A Matlab Toolbox for Circular Statistics, Journal of Statistical Software, Volume 31, Issue 10, 2009, http://www.jstatsoft.org/v31/i10), for each frequency and target position. A Harrison-Kanji test was performed (two-way ANOVA for circular data; circ_hktest from the Circular Statistics Toolbox) on the individual optimal phases, with frequency and position as factors. An interaction would indicate that the effect of position on the optimal phase differs across frequency conditions. We further computed the pairwise phase difference with the function circ_dist participant-by-participant, for each pair of positions (positions 1 and 2, 1 and 3, 2 and 3), and we tested whether the pairwise phase difference was significantly different against 0, i.e., null difference, for each frequency and pair of positions, with a Watson-Williams test (function circ_wwtest) from the Circular Statistics Toolbox. P-value were Bonferroni corrected for multiple comparisons across target positions and frequencies (p-value threshold < 0.004, corresponding to a Z-score threshold of 2.64, and a Cohen’s d threshold of 1.66).

### Control experiment 1: Flashing disk

This control was designed based on Sokoliuk and VanRullen (2016).

#### Experimental Design

The targets were of same size and location as in the main experiment. The peripheral disk was, however, not modulated sinusoidally but was flashed (during three frames, for a total of 6.3 ms) at 7 different levels of luminance (7 contrast levels: 5%, 20.8%, 36.7%, 52.5%, 68.3%, 84.2% and 100%) at the same time as target onset to test for luminance masking. There was 20% of catch trials in which the disk was flashed but not the target.

As for the main experiment, the control experiment first included a staircase procedure to adjust the contrast of each target, for each participant, to reach 50% detection. In 7 blocks, the luminance of the disk was set at 50% contrast. For each block, 6 to 18 targets were presented (∼10 targets/trials per block on average), thus the staircase procedure contained an average of 70 trials.

Then, participants performed four runs of 38 blocks each in two separate experimental sessions of approximately 1h30 each (no EEG recording for this control experiment). Each contrast level was displayed the same number of times (∼217 targets per contrast level). Target contrasts were adjusted every 15 blocks to maintain the same detection level across the entire session.

#### Analyses

Hit rates, i.e., percentage of correct detection, median reaction times and target contrasts were computed for each target position. Hit rates were 57.18 ± 1.1 % (mean ± std), 51.74 ± 0.35 %, and 45.82 ± 0.76 % for position 1, 2 and 3, respectively. Median reaction times were 474 ± 2 ms (mean ± std), 511 ± 3 ms, and 525 ± 3 ms, for position 1, 2, and 3, respectively. Target contrasts were 0.61 ± 0.01 % (mean ± std), 1.16 ± 0.04 %, and 0.71 ± 0.02 %, for position 1, 2, and 3, respectively. A one-way repeated-measures ANOVA was performed for each dependent variable to assess the effect of target position. For hit rates, there was no main effect of target position (F(1,14) = 2.52, p-value = 0.09, eta² = 15.29). For median reaction times, there was also no main effect of target position (F(1,14) = 1.24, p-value = 0.30, eta² = 8.16). For target contrast, there was a main effect of target position (F(1,14) = 13.68, p-value < 0.01, eta² = 49.42), which is coherent with the staircase manipulation.

#### Luminance binning

Targets were binned according to the level of luminance of the simultaneously presented disk. To emulate an oscillatory cycle (which, in the main experiment, goes from 0% contrast to 100% and back to 0%), we used a bootstrap procedure (5 000 repetitions) randomly assigning half of the targets to one half of the cycle (from contrast 0% to 100%), while the other half was assigned to the second half of the cycle (from contrast 100% to 0%), thus resulting in 13 luminance levels.

The number of false alarms (i.e., participants responded a target was present while it was absent), and the number of hits were computed for each target position and for the 13 luminance levels. A one-cycle sine function was fitted to individual data and data averaged across participants (Eq2) with the following free parameters: the amplitude, *x*[1], the phase offset, *x*[2], the baseline, *x*[3], and a Least Squares cost function was used to find the parameters that best fit the data.

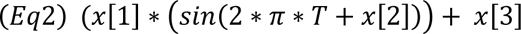

We performed a Monte Carlo procedure on the fitted amplitudes to assess whether the fitted data was modulated periodically by the luminance disk (see **Statistical analyses** section). There was no significant effect of luminance, at any target position, for the false alarm ratio and thus we did not consider this variable any further.

The optimal phase, i.e., phase of highest performance, was computed for each target position on data averaged across participants. To study whether the optimal phase shifted as a function of target positions, we compared the individual optimal phase and assessed the significance using a one-way repeated measures ANOVA (main variable: individual optimal phase, main factor: target positions).

### Control experiment 2: Broadband oscillating disk

#### Experimental Design

Targets were of same size and location as in the main experiment and control experiment 1. The peripheral disk was, however, oscillating in luminance with a broadband frequency profile centered on 10 Hz (from 4.1 Hz to 23.9 Hz, excluding 10 Hz). This manipulation allowed testing potential masking effects under the same attentional condition as in the main experiment (note that in the previous control, the flashing disk could have attracted attention in the periphery). Following the same staircase procedure as in the main experiment, participants then performed three runs of 42 blocks each, over two separate experimental sessions of ∼1h30 each. In one of the two sessions of two of the participants, EEG was simultaneously recorded (same analysis as in the main experiment, see **EEG Analysis** above).

#### Analyses

Hit rates, median reaction times and target contrasts were computed for each target position. Hit rates were 46.1 ± 1.11 % (mean ± std), 44.68 ± 1.22 %, and 46.9 ± 0.96 % for position 1, 2 and 3, respectively. Median reaction times were 515 ± 9 ms (median ± std), 520 ± 9 ms, and 515 ± 8 ms, for position 1, 2, and 3, respectively. Target contrasts were 3.7 ± 0.55 % (mean ± std), 2.2 ± 0.26 %, 2.29 ± 0.28 %, for position 1, 2, and 3, respectively. A one-way repeated-measures ANOVA was performed for each dependent variable to assess the effect of target position. There was no main effect of target position for both hit rates (F(1,14) = 3.45, p-value = 0.05, eta² = 19.8), and median reaction times (F(1,14) = 1.13, p-value = 0.34, eta² = 7.44). For target contrast, there was a main effect of target position (F(1,14) = 23.95, p-value < 0.01, eta² = 63.11), which is coherent with the staircase manipulation.

#### Luminance binning

Data were analyzed following the same procedure described in the **Control experiment 1: Flashing disk** (see *Luminance binning* section). Targets were binned according to the level of luminance of the oscillating disk, i.e., 7 equally spaced bins from 0% to 100% levels of contrast followed by a bootstrap procedure to obtain 13 luminance levels. A one-cycle sine function was fitted to individual data and data averaged across participants (Eq2), and a Least Squares cost function was used to find the parameters that best fit the data. A Monte Carlo procedure on the fitted amplitudes was performed to assess whether the fitted data was modulated periodically by the luminance disk. Differences in optimal phase across target position was tested with a one-way repeated measures ANOVA (main variable: individual optimal phase, main factor: target positions).

#### Luminance peak binning

To assess whether the maximal luminance of the peripheral disk could induce behavioral masking which would then propagate across the retinotopic space, targets were assigned to time bins of a 100 ms periods following the peaks in disk luminance. Here again, a one-cycle sine function (Eq2) was fitted to the data and a Monte Carlo procedure assessed significance. Differences in optimal phase across target position was tested with a one-way repeated measures ANOVA (main variable: individual optimal phase, main factor: target positions).

#### Pseudo-phase binning

To assess whether the arrhythmic, broadband (around, but not including, 10 Hz) peripheral disk produced the same results as in the main experiment in which the disk is oscillating at 10 Hz, targets were assigned to phase bins of 100 ms periods starting from disk onset. As before, a one-cycle sine function was fitted to individual data and data averaged across participants (Eq2) and a Monte Carlo procedure on the fitted amplitudes was performed to test whether the data was modulated at 10 Hz.

### Statistical Analyses

To test whether the fitted data was significantly modulated periodically by the phase of the oscillating disk (see **Phase effect on detection performance** section), a Monte Carlo procedure on the fitted amplitude was performed. 50,000 surrogates were generated for each participant, target position and frequency, by randomly assigning correct responses, i.e., for each target, we randomly assigned an incorrect or a correct response based on the averaged performance, and false alarms, i.e., a number of false alarms at an associated random delay was randomly assigned to each block, based on the average number of false alarms throughout the experiment. Hit and false alarm rates were then computed for each phase bin, position, and frequency, and the same sine function were fitted (Eq1). The surrogate distributions of the 50,000 fitted amplitudes for the induced frequency (free parameter x[1] from (Eq1)) and the first harmonic (free parameter x[3] from (Eq1)) were compared to the empirical fitted amplitudes, i.e., the surrogate fitted amplitudes of each component (induced, first harmonic, sum of both) were respectively compared to the empirical fitted amplitudes. P-values were obtained by computing the proportion of fitted amplitudes equal or higher than the empirical fitted amplitude (fitted amplitudes: free parameters *x*[1] and *x*[3] from (Eq1)). The fitted amplitudes of the induced frequency (*x*[1]) and of the first harmonic (*x*[3]) across all frequencies and target positions were compared with two-tailed t-test to investigate which frequency contribute the most to the oscillatory pattern.

In the control experiments (see **Control experiment** section), the same Bonferroni corrected Monte Carlo procedures were performed, except that the 50,000 datasets were created for each binning conditions and that the sine function (Eq2) was fitted to the data.

## Results

Participants (n=17) performed a threshold visual detection task, while a luminance oscillating disk was concurrently presented in the periphery (eccentricity: 7.5°) to modulate perception at the theta and alpha frequency (4, 6, 8 and 10 Hz; EEG simultaneously recorded). Near-threshold target stimuli appeared at one of three possible eccentricities between a central fixation cross and the disk (**Figure 1A**; adjusted according to cortical magnification so each target measured 0.8 mm of diameter and were placed 0.8 mm away from each other in the cortex; see **Figure 1B**). We tested whether (1) the periodic disk stimulation modulates detection performance periodically at each target position, (2) the optimal phase (of highest performance) shifts as a function of distance from the disk suggesting that induced perceptual cycles travel across space (**Figure 1C**), and (3) the occurrence of such traveling properties depends on the induced frequency.

**Figure 1.**
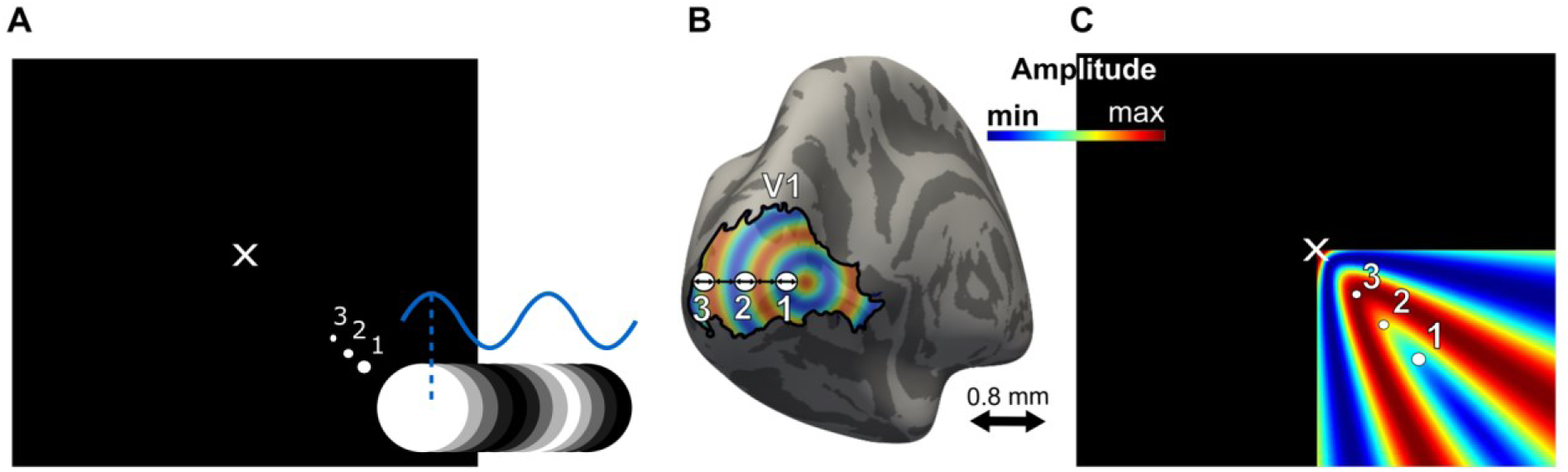
Experimental protocol. **A.** A luminance oscillating disk (inducer) was presented in the periphery to induce perceptual cycles at the theta and alpha frequency. Participants were instructed to detect visual targets at threshold (50% detection) at three different positions in the retinotopic space, based on cortical magnification. Disks oscillated for a 30-seconds period during which 6 to 18 targets were presented at random times according to a decreasing probability distribution. Participants pressed the space bar when they detected the target (1-s response window after target onset) **B.** In the visual cortex, visual targets measured 0.8 mm of diameter and were placed 0.8 mm away from each other. **C.** Hypothesis: the oscillating disk, presented in the lower right corner, induced a brain activity that traveled across the retinotopic space (note that we concentrate our predictions to the quadrant in which the disk is presented). At an instant t, position 3 is located at the maximum amplitude of the induced brain activity, leading to the highest performance, while position 1 is located at the minimum amplitude.

EEG activity was analyzed using frequency decomposition (Fast Fourier Transform, FFT, performed on the time series of each participant and electrode) and Event-Related Potentials (ERPs) measures to ensure that we successfully induced frequency-specific brain activity and to characterize the shape of the evoked response. First, peaks in the spectrum (as measured per SNR, averaged across participants and electrodes) were identified for each induced frequency (4, 6, 8 and 10 Hz; SNR > 1.4) and their first harmonics (8, 12, 16 and 20 Hz; SNR > 1.57), with topographies showing brain activity in the occipital cortex (for simplicity, frequencies were rounded to a whole number; real values are displayed in **Figure 2 left side of each panel**). Second, ERP analyses showed that the evoked signal is a complex signal composed of the induced frequency and its first harmonic (**Figure 2 right side of each panel**). We further computed an FFT on the ERPs of electrode Oz. Peaks were again identified for each induced frequency (4, 6, 8 and 10 Hz; SNR > 2.4) and their first harmonics (8, 12, 16 and 20 Hz; SNR > 4.41; data not shown). Note that both SNR computed on EEG time series and on EEG evoked activity are > 1 for each induced frequency and first harmonic demonstrating that the disk induced frequency-specific brain activity. These complex evoked responses were interpreted as the overlap between the periodic brain response (i.e., inducer) and the neural population response to individual stimuli (i.e., contrast change) at low frequencies, or alternatively, to the nonlinear nature of the visual system, i.e., nonlinear systems produce complex output consisting of the input frequency and multiple harmonics (Heinrich, 2010; Norcia et al., 2015). Together, the EEG analyses confirm that we successfully induced theta/alpha activity in the visual cortex and show that the evoked signal is a complex signal composed of the induced frequency and its first harmonic (Heinrich, 2010; Norcia et al., 2015).

**Figure 2.**
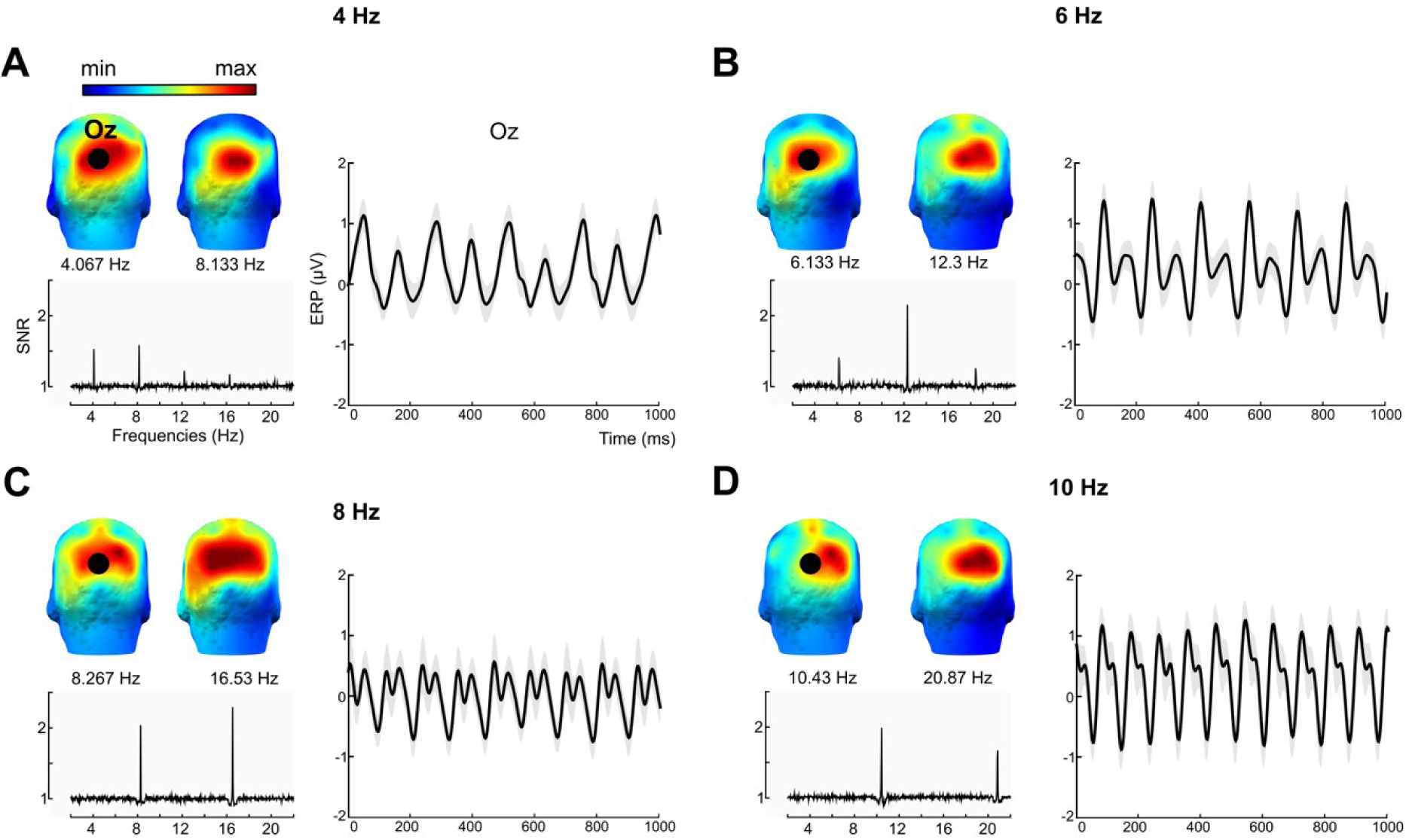
The oscillating disk induced frequency-specific brain activity in the occipital cortex. Left side of each panel, Fast Fourier Transform analyses performed on the EEG time series. Right side of each panel, ERP analyses, for induced frequencies of **A.** 4, **B.** 6, **C.** 8 and **D.** 10 Hz (rounded frequency values). Signal-to-noise ratio (SNR) amplitude averaged across 17 participants and all 62 electrodes, from 2 to 22 Hz. In each panel, a peak in SNR is identified for the induced frequency and its first harmonic. Topographies of the two SNR peaks revealed that we successfully induced brain activity in the visual cortex. Black dot, Electrode Oz. ERP of the Oz electrode computed on one-second epochs (the first-second epoch was removed to each 30-seconds blocks to avoid the EEG transient evoked response; each block represented 29 1-second epochs) averaged across participants (n=17). Solid black line, averaged ERP. Shaded area, standard error of the mean.

Second, we investigated whether detection performance was modulated periodically for each induced frequency and each target position. Targets (from 6 to 18 targets per 30 seconds-blocks) can appear at random delays during the periodic disk stimulation (from 1s to 29s from disk onset). Targets were binned according to the phase of the oscillating disk (7 bins/cycle; least number of bins to sample one oscillatory period while allowing a large number of trials per bin to achieve high statistical power). In each bin, detection performance was computed as per hit rate (correct target detection) for each participant, frequency, and target position. Finally, the data were averaged across participants and given the complex shape of the ERP (**Figure 2**), fit to a complex sine function (the shape of the evoked response was used as proxy for the shape of the behavioral response), separately for each frequency and target position. Detection performance showed a significant oscillatory pattern for each target position and frequency (Monte Carlo on fitted amplitude, Bonferroni corrected across frequencies and positions, p-value threshold < 0.004, corresponding to a Z-score threshold of 2.64, and a Cohen’s d threshold of 1.66; **Figure 4**), and a clear optimal phase (of highest performance). Note that the fitted amplitude of the induced frequency (distance between the optimal and non-optimal phase) was significantly higher than the one of the first harmonic (two-tailed t-test on fitted amplitude across all frequencies and positions, p-value < 0.01, Cohen’s d = 0.67, CI = [0.05; 0.1]). A high fitted amplitude indicates that the fitted sine function explains a large portion of the variance in the behavioral performance. The results thus suggest that the oscillatory pattern was mainly driven by the induced frequency across all tested target positions and frequencies. The inducer modulated detection performance periodically, at each target position and at each frequency, with an average amplitude modulation (optimal to non-optimal hit rate difference) of 42.71 ± 3.37 %. The amplitude modulation was further computed for each participant, frequency, and target position (**Figure 3**). A two-way repeated-measures ANOVA revealed a significant main effect of the frequency (F(3, 48) = 23.85, SS = 0.67, p-value < 0.01, eta² = 59.85), and no main effect of the target position (F(2, 32) = 0.9, SS = 0.01, p-value = 0.41, eta² = 5.33), nor of the interaction (F(6, 96) = 0.92, SS = 0.03, p-value = 0.48). The amplitude modulation was of 49.84 ± 3.45% for 4 Hz, 46.56 ± 3.57 % for 6 Hz, 39.04 ± 2.67 % for 8 Hz, and 35.39 ± 2.54 % for 10 Hz. Post-hoc two-tailed t-tests showed that the amplitude modulation was higher for 4 Hz compared to 8 Hz (p-value < 0.01, Cohen’s d = 0.84, CI = [0.07; 0.14]) and 10 Hz (p-value < 0.01, Cohen’s d = 1.15, CI = [0.1; 0.18]), with a tendency compared to 6 Hz (p-value = 0.06, Cohen’s d = 0.22, CI = [-0.01; 0.06]), higher for 6 Hz compared to 8 Hz (p-value < 0.01, Cohen’s d = 0.57, CI = [0.04; 0.1]) and 10 Hz (p-value < 0.01, Cohen’s d = 0.87, CI = [0.07; 0.14]), and higher for 8 Hz compared to 10 Hz (p-value = 0.02, Cohen’s d = 0.33, CI = [0.01; 0.06]). In summary, the amplitude modulation decreases with increasing frequency but is constant across target positions. The amplitude modulation allowed to reliably estimate the optimal phase at each frequency and target positions. A complementary analysis showed that behavioral performance was better explained when taking into account the spatial position of the targets. Specifically, we compared the amplitude modulation when detection performance was calculated disregarding target position, with the amplitude modulation calculated independently for each target position and then averaged across positions (two-tailed t-tests Bonferroni corrected for multiple comparisons: 4 Hz, t-stat = -5.84, p-value < 0.01, Cohen’s d = -0.3, CI = [-0.05; -0.02]; 6 Hz, t-stat = -5.45, p-value < 0.01, Cohen’s d = -0.25, CI = [-0.04; -0.02]; 8 Hz, t-stat = -5.35, p-value < 0.01, Cohen’s d = -0.3, CI = [-0.04, -0.01]; 10 Hz, t-stat = -9.1, p-value < 0.01, Cohen’s d = -0.52, CI = [-0.05; -0.03]). The amplitude modulation was significantly stronger when calculated position-by-position, suggesting that taking into account the spatial position of visual targets better explain perceptual performance.

**Figure 3.**
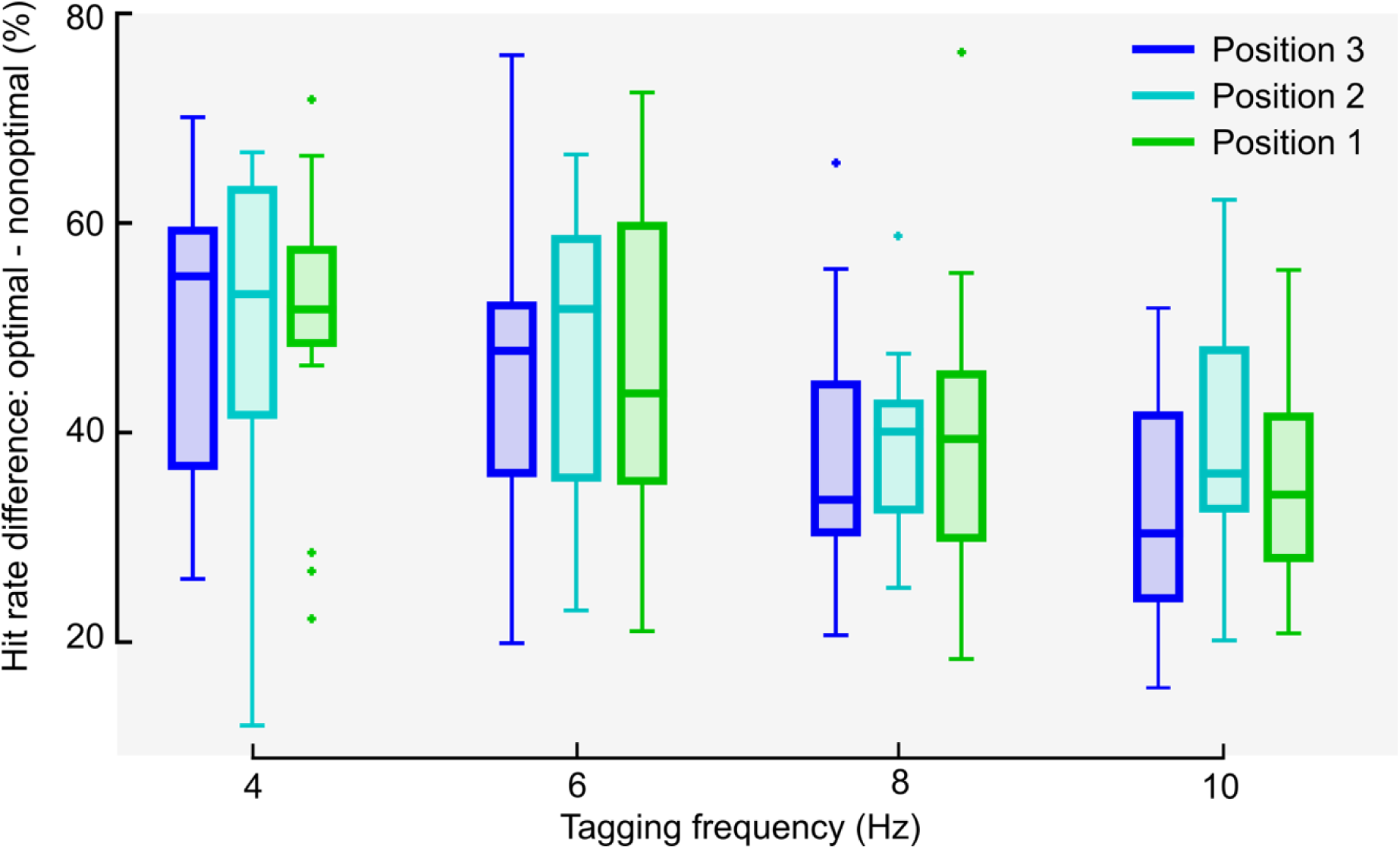
The amplitude modulation decreased with increasing induced frequency. Amplitude computed as the difference between the maximum and the minimum hit rate. Dots, individual outlier data. A two-way repeated-measures ANOVA revealed a significant main effect of frequency (F(3, 48) = 23.85, SS = 0.67, p-value < 0.01, eta² = 59.85).

Critically, we tested whether (1) perceptual cycles propagate across the retinotopic space, and (2) the occurrence of such traveling properties depends on the induced frequency. Propagation is defined as a shift in the optimal phase between two positions. To assess whether the optimal phase shifted as a function of target position, individual optimal phases were converted in radians based on a one-cycle sine wave fit (see **Methods**) at the induced frequency (the oscillatory pattern was mainly driven by the induced frequency; see analysis above). A Harrison-Kanji test revealed a significant main effect of the frequency (F(3,48) = 25.13, SS = 14.33, p-value < 0.01), target position (F(2,32) = 3.06, SS = 1.16, p-value = 0.04) and a tendency for the interaction (F(6,96) = 1.99, SS = 2.42, p-value = 0.06). For the post-hoc analyses, we performed Watson-Williams tests against 0, i.e., null difference, on participant-by-participant pairwise phase differences, for each pair of positions (positions 1 and 2, 1 and 3, 2 and 3). The results showed a significant phase shift between positions 1 and 2, 1 and 3, and 2 and 3, for both 8 Hz and 10 Hz (**Figure 4C,D**; Bonferroni corrected across frequencies and positions, p-value < 0.004); and between positions 1 and 2 for 6 Hz. Thus, we did not find a systematic phase shift, i.e., phase shift between *each* target positions, for 4 and 6 Hz (**Figure 4A,B**). Together, the results demonstrate that the behavioral optimal phase of alpha perceptual cycles shifted across target positions, suggesting that brain signals traveled across the retinotopic cortical space. In addition, the phase shift cannot be explained by a decreased phase estimation accuracy at low frequencies (see **Figure 3**, showing that the amplitude modulation is in fact higher at low compared to high induced frequency).

**Figure 4.**
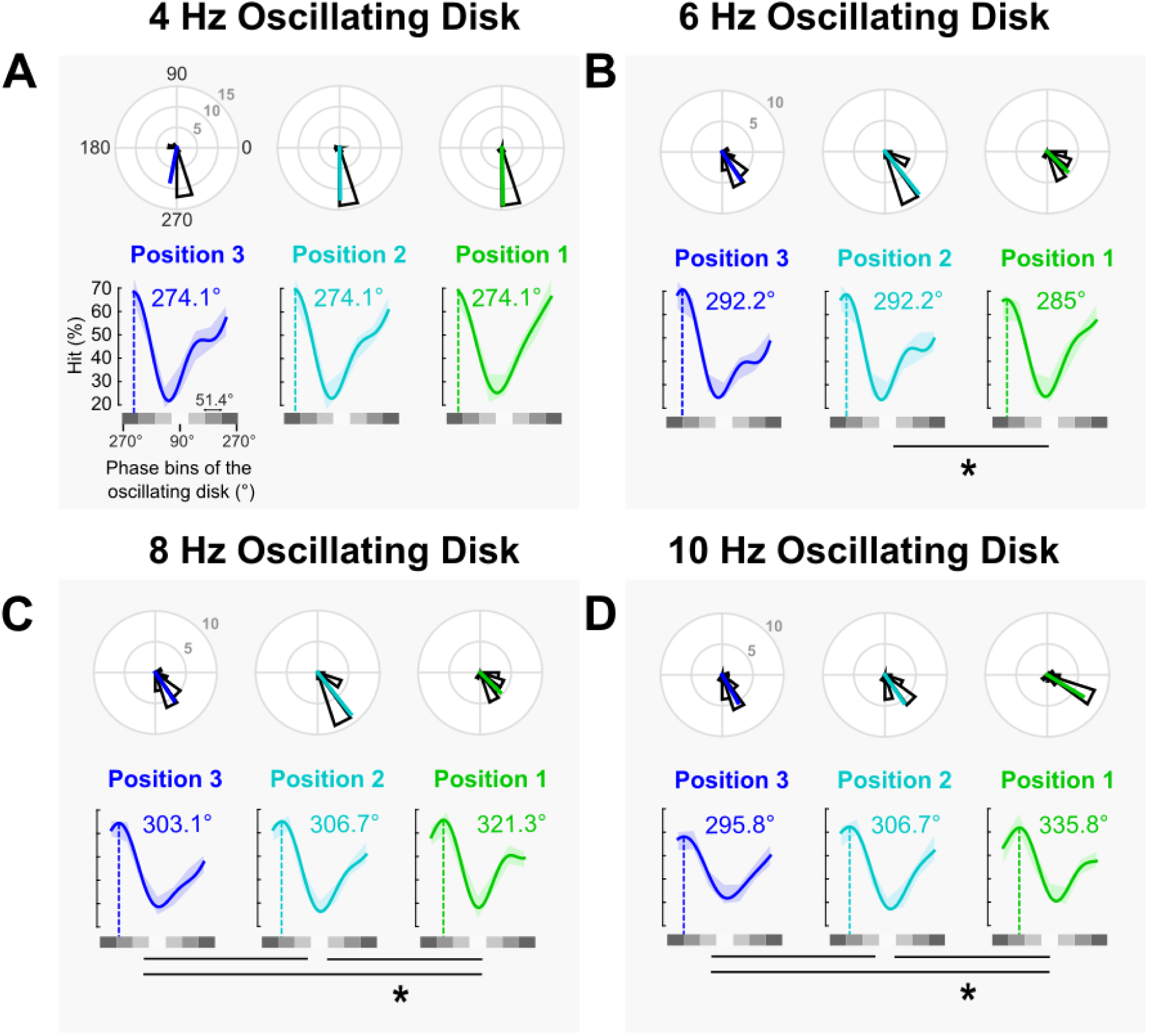
The optimal phase of alpha perceptual cycles shifts between target positions. Lower graph of each panel, hit rate averaged across participants (n=17), binned as a function of the phase of the oscillating disk, and fitted to a complex sine function. Shaded colored area, 95 confidence intervals. Solid line, sine fit. Dotted line, optimal phase in degrees. The value of the optimal phase in degrees is extracted for fit on data averaged across participants (note that this closely resemble the value obtained from averaging individual optimal phases from top graph). Top graph of each panel, rose plot distribution of the optimal phase across participants. Colored solid line, optimal phase averaged across participants. Gray scale, phase bins of the oscillating disk (90°, white disk). Each bin represents 51.4° of the oscillatory cycle. **A.** results for induced frequencies of **A.** 4, **B.** 6, **C.** 8 and **D.** 10 Hz. Performance was significantly modulated periodically at each target position and frequency (Monte Carlo, Bonferroni corrected across target positions and frequencies, p-value threshold < 0.01). The optimal phase shifted between positions 1 and 2, 2 and 3, and 1 and 3 for 8 Hz- and 10 Hz-induced frequencies; and between positions 1 and 2 for 6 Hz-induced frequency (Watson-William test, Bonferroni corrected across target positions and frequencies, p-value threshold < 0.004). There was no significant phase shift for 4 Hz-induced frequencies.

When converting the phase shift observed between target positions into milliseconds (3 ms for 8 Hz, and 5.5 ms for 10 Hz, respectively) and given the cortical distance between two target positions (0.8 mm distance edge-to-edge) as well as the targets’ cortical size (0.8 mm diameter), one can estimate that alpha-induced brain oscillations traveled across the retinotopic space at an averaged propagation speed ranging between 0.3 (1.6 mm / 5.5 ms) and 0.5 m/s (1.6 mm / 3 ms).

Finally, one may ask whether the phase shift observed in behavioral data was due to luminance masking from the disk. We performed two control experiments (15 participants each) to address this potential concern. Participants were instructed to detect visual targets at threshold (50% detection) at one of the same three positions as in the main experiment, while a peripheral disk was simultaneously flashed at one of seven possible levels of luminance (**Control Experiment 1: Flashing Disk**; same as Sokoliuk and VanRullen, 2016), or while the disk was oscillating in luminance with an arrhythmic broadband frequency profile centered on 10 Hz (**Control Experiment 2: Broadband Oscillating Disk**). First, for both control experiments, detection performance was calculated for each level of luminance, and a one-cycle sine function was fitted to the empirical data (see **Luminance binning** in **Methods**). For both control conditions, detection performance showed a significant oscillatory pattern at each target position (Monte Carlo, Bonferroni corrected across positions, p-value threshold of 0.01, corresponding to a Z-score threshold of 2.12, and a Cohen’s d threshold of 1.31, **Figure 5A,B**) confirming the presence of luminance masking at all three positions. We then compared the individual optimal phase between target positions. There was no significant phase shift in both the Flashing Disk control (F(2, 14) = 1, SS = 345.5, p-value = 0.38, eta² = 6.67) and the Broadband Oscillating Disk control (F(2, 14) = 0, SS = 3.61, p-value = 0.99, eta² = 0), suggesting that the propagation of perceptual cycles observed in the main experiment was not due to masking.

**Figure 5.**
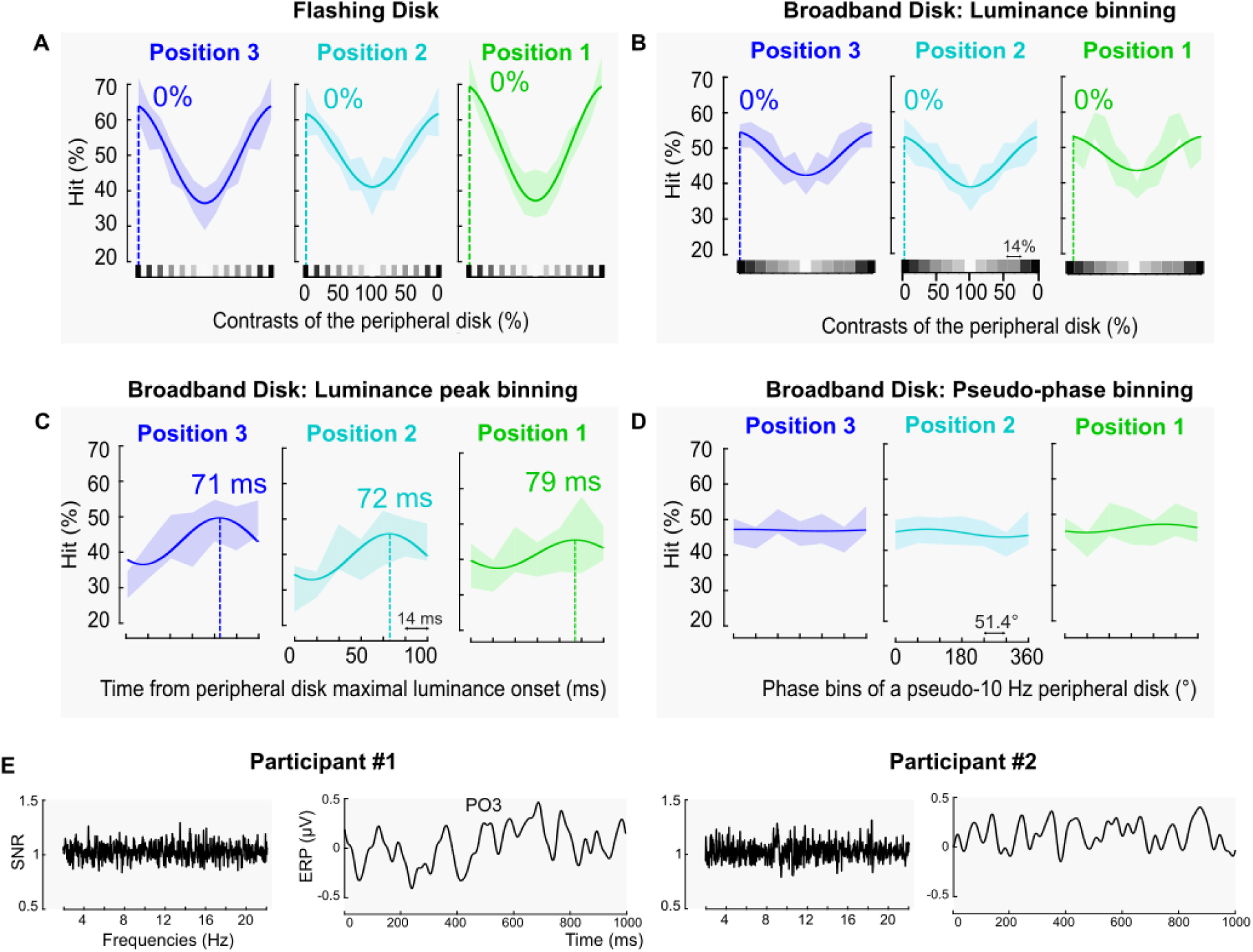
The propagation of induced perceptual cycles was not due to luminance masking. **A.** Flashing Disk control: Hit rate for each 13 discrete luminance levels and 3 target positions averaged across participants (n=15) and fitted to a one-cycle sine function. Shaded color area, 95 confidence intervals. Solid line, fit. Dotted line, optimal behavioral phase. Green, position 1. Cyan, position 2. Blue, position 3. **B.** Broadband Oscillatory Disk control: Hit rate for each 13 luminance bins and 3 target positions averaged across participants (n=15) and fitted to a one-cycle sine function. **C.** Hit rate for each 7-time bins from the peripheral disk maximal luminance onset, and the 3 target positions averaged across participants (n=15) and fitted to a one-cycle sine function. **D.** Hit rate for each 7 pseudo-phase bins of 10 Hz rhythmic modulation, and the 3 target positions averaged across participants (n=15) and fitted to a one-cycle sine function. **E.** EEG data of two individual participants. Left side of each panel, SNR amplitude averaged across all 62 electrodes on the Fast Fourier Transform performed on the EEG time series. Right side, ERP of the PO3 electrode computed on one-second epochs (the first-second epoch was removed to each 30-seconds blocks to avoid the EEG transient evoked response; each block represented 29 1-second epochs).

To further investigate whether the maximal luminance of the disk (100 % contrast) could induce a propagation of behavioral masking, targets in the Broadband Oscillating Disk Experiment were binned as a function of the peak luminance (see **Luminance peak binning** in **Methods**). Visual performance showed a significant oscillatory modulation for position 1 and 2 (Monte Carlo, Bonferroni corrected across positions, p-value threshold of 0.01, corresponding to a Z-score threshold of 2.12, and a Cohen’s d threshold of 1.31) but not position 3 (p = 0.04, above the Bonferroni corrected p-value threshold) (**Figure 5C**). Although the fluctuation was not significant at position 3, we still compared the individual optimal phases between target positions. The test did not reveal a significant phase shift (F(2, 14) = 0.7, SS = 590.93, p-value = 0.5, eta² = 4.82). Coherent with the previous result, although the disk maximal luminance masked visual performance of nearby targets, this masking cannot explain the shift of the optimal phase in the main experiment.

We finally investigated whether the arrhythmic, broadband peripheral disk produced the same results as in the main experiment by binning the targets every 100 ms (see **Pseudo-phase binning** in **Methods**). Although centered at 10 Hz, the broadband oscillating disk did not periodically modulate detection performance at 10 Hz (Monte Carlo, p-value > 0.44, **Figure 5D**). This absence of rhythmic modulation was coherent with the EEG analysis on two participants showing that the broadband peripheral disk did not evoke a frequency-specific 10 Hz brain activity (see SNR results on **Figure 5E**; although it did induce broadband activity around 10 Hz, analysis not shown). In other words, the broadband visual stimulation did not induce narrow band 10 Hz-brain activity, coherent with the behavioral results showing no alpha induced perceptual cycle.

Taken together, the results of both control experiments show that the phase shift observed in the main experiment cannot be merely explained by low-level luminance masking (**Figure 5**).

## Discussion

Using psychophysics and EEG, we showed that visual perception at each target position is modulated periodically at the induced frequency, replicating previous results (Stefanics et al., 2010; Mathewson et al., 2012; Henry and Obleser, 2012; Graaf et al., 2013; Henry et al., 2014; Spaak et al., 2014; ten Oever and Sack, 2015; Kizuk and Mathewson, 2017). Critically, the optimal phase for visual perception systematically shifted between target position. This effect was observed for alpha-induced frequencies (8 Hz and 10 Hz; and cannot be simply explained by a mere luminance masking confound) and not theta, with an average propagation speed ranging from 0.3 to 0.5 m/s. Together, the results support the hypothesis that induced perceptual cycles propagate across the retinotopic visual space presumably due to horizontal connections in the retinotopic visual cortex.

### Entrained oscillations or periodic evoked response?

Our results showed that perceptual cycles are propagating across the *visual* retinotopic space, suggesting that the underlying neural activity is propagating as well in the *cortical* retinotopic space through horizontal axons. We cannot, however, conclude whether this propagating activity corresponds to the propagation of *true* entrained oscillations or of a succession of evoked responses. Our EEG recordings highlighted that the evoked signal was composed of the induced frequency and its first harmonic, interpreted as the overlap between the periodic brain response (i.e., inducer) and the neural population response to individual stimuli (i.e., contrast change) at low frequencies, or alternatively, to the nonlinear nature of the visual system, i.e., nonlinear systems produce complex output consisting of the input frequency and multiple harmonics (Heinrich, 2010; Norcia et al., 2015). The propagation we observed in this study being frequency band-specific, we argue that the most parsimonious interpretation is to consider entrained brain oscillations. However, and regardless the underlying cortical mechanism, whether entrained oscillations or periodic ERPs, our conclusions remain unchanged, i.e., induced perceptual cycles propagate across the retinotopic space.

### Propagation of alpha but not theta perceptual cycles

Attention is the cognitive function that selects relevant information to facilitate its processing while still being able to flexibly redeploy resources to other information if necessary (Carrasco, 2011; Dugué et al., 2018, 2020). Studies have shown that when covert attention (no head/eye movements) is sustained at a given spatial location, information is sampled periodically at the alpha frequency (review: Dugué and VanRullen, 2017; Kienitz et al., 2022). In other words, visual performance fluctuates over time in the alpha band in detection tasks in which the target stimulus appeared always at the same spatial location (Valera et al., 1981; Busch et al., 2009; Mathewson et al., 2009; Busch and VanRullen, 2010; Dugué et al., 2011; Samaha and Postle, 2015; Samaha et al., 2017; Fakche et al., 2022). However, when multiple stimuli are presented, attention rhythmically samples information at the theta frequency (Dugué et al., 2015; Landau et al., 2012; Fiebelkorn et al., 2013a; Song et al., 2014; Huang et al., 2015; Dugué et al., 2019; Merholz et al., 2022; Galas et al., 2023; see also Re et al., 2023; review: VanRullen, 2016; Dugué and VanRullen, 2017; Kienitz et al., 2022; Keitel et al., 2022).

Here, while we did not explicitly manipulate covert attention, participants fixated at the center of the screen and targets appeared in a constrained location suggesting that covert, voluntary spatial attention was sustained on the bottom right quadrant. This manipulation likely recruited alpha sampling, which potentially favored the propagation of alpha-induced perceptual cycles. One can speculate that the propagation of alpha perceptual cycles away from the main attention focus could allow the observer to periodically monitor other nearby locations allowing for flexible attentional reallocation when a target appears. Further studies are necessary to investigate this hypothesis.

### Perceptual propagation suggests cortical propagation

Sokoliuk and VanRullen (2016) reasoned that if an induced brain activity propagates across the visual cortex, it should have perceptual consequences across the retinotopic visual space. Using a similar psychophysical paradigm, they found a shift of the behavioral optimal phase as a function of distance from the inducer. Yet, there are two notable differences between their study and the present one. First, they observed that the phase shifted between the visual stimulus the closest to the inducer and more distant ones (no differences between intermediate positions). Yet, postulating brain activity is traveling within the cortex requires showing a phase change across all positions. Here, contrary to Sokoliuk and VanRullen (2016), we found a systematic phase shift across all three positions.

Second, Sokoliuk and VanRullen (2016) observed an optimal phase shift for both induced-frequencies of 5 Hz (theta) and 10 Hz (alpha), but not 15 Hz. Here, however, we showed a shift for alpha-induced frequencies (8 Hz and 10 Hz), but not theta (4 Hz and 6 Hz).

Together, these discrepant results can be explained by a number of critical differences between the two studies. First, the EEG analysis confirmed that the oscillating disk induced frequency-specific brain activity in the occipital cortex. This result was important to validate our rationale postulating that induced perceptual cycles travel across the *visual* retinotopic space due to the propagation of brain activity across the *cortical* retinotopic space. Importantly, our EEG results revealed that the oscillating disk produced a complex neural response composed of the induced frequency and its first harmonic, coherent with the nature of the visual system (Heinrich et al., 2010; Norcia et al., 2015; Spaak et al., 2014). Thus, one must fit the behavioral response with a corresponding complex sine function to accurately capture the non-linearity of the neural response. Since the EEG was not recorded in Sokoliuk and VanRullen (2016), the shape of the induced brain signal was unknown. Second, to ensure that the three targets activated the same number neurons in visual regions, hence landing in a similar spatial extent (spatial phase) of the propagating activity, we adjusted the size of each target to cortical magnification. Third, in our study, the same participants (n=17) performed all four frequency conditions thus ensuring equal statistical power and no inter-individual variability (n=5 for 5Hz, n=7 for 10Hz and n=15 for a 10-Hz replication set, and n=4 for 15Hz in Sokoliuk and VanRullen, 2016). Finally, we used eye-tracking to ensure stable fixation, critical when investigating retinotopic propagation.

### Neural mesoscopic traveling waves, a growing, yet largely understudied field

A traveling wave is the propagation of neural activity over space with a monotonic shift in the peak latency between the origin of the signal and more distal positions (Sato et al., 2012; Muller et al., 2018).

Mesoscopic waves travel *within* individual brain regions (e.g., V1) spanning millimeters (Sato et al., 2012; Muller et al., 2018). Studies using invasive recordings with high spatial and temporal resolution showed non-oscillatory traveling activity in visual cortices of mammals (Bringuier et al., 1999; Jancke et al., 2004; Roland et al., 2006; Lippert et al., 2007; Sharon et al., 2007; Xu et al., 2007; Takagaki et al., 2008; Nauhaus et al., 2009, 2012; Gao et al., 2012; Rekauzke et al., 2016; Slovin et al., 2002; Chen et al., 2006; Sit et al., 2009; Reynaud et al., 2012; Zhang et al., 2012; Yang et al., 2015; Chemla et al., 2019). Much fewer studies have focused on mesoscopic *oscillatory* traveling waves (spontaneous, entrained or evoked). They showed that delta, theta, alpha, beta and gamma oscillations can propagate within individual visual areas of mammals and turtles (Prechtl et al., 1997, 2000; Sanchez-Vives and McCormick, 2000; Huang et al., 2004; Benucci et al., 2007; Han et al., 2008; Ray and Maunsell, 2011; Maris et al., 2013; Stroh et al., 2013; Muller et al., 2014; Townsend et al., 2015; Zanos et al., 2015; Davis et al., 2020). In humans, only a few studies, using invasive recordings in patients, showed the propagation of theta, alpha, and beta mesoscopic traveling waves (Takahashi et al., 2011; Zhang and Jacobs, 2015; Sreekumar et al., 2021), but only in non-visual cortical areas. There is no electro-physiological study to date investigating the propagation of low-frequency oscillations in individual human visual areas (see Grabot et al., 2022, for a computational solution).

Finally, if a few studies have investigated the functional role of propagating oscillatory activity at the mesoscopic levels (attention: Maris et al., 2013; memory: Han et al., 2008; motricity: Rubino et al., 2006; Takahashi et al., 2011, 2015; saccadic eye movements: Zanos et al., 2015; visual perception: Besserve et al., 2015; Davis et al., 2020), very little is known about the link between oscillatory activity propagation within individual visual areas and perceptual performance, especially in human. The present study addresses this clear gap in the literature.

### Speed and spatial extent of mesoscopic traveling waves

Our results show that induced alpha perceptual cycles travel across the retinotopic space at a propagation speed ranging from 0.3 to 0.5 m/s. Such observation is in line with results from the animal literature showing that low-frequency oscillations propagate within visual areas at a speed ranging from 0.01 to 0.6 m/s (Prechtl et al., 1997, 2000; Sanchez-Vives and McCormick, 2000; Benucci et al., 2007; Han et al., 2008; Ray and Maunsell, 2011; Muller et al., 2014; Zanos et al., 2015; Davis et al., 2020). Additionally, given a phase difference of maximum 40° between target positions, it is unlikely that more than one (spatial) cycle (360°) of oscillatory activity propagates across such a small portion of the visual cortex, in line with previous observations (Zanos et al., 2015; Zhang and Jacobs, 2015).

Finally, given our specific contrast manipulation and the observed propagation of induced perceptual cycles, it is likely that our paradigm preferentially probed alpha traveling waves in area V1 (with small receptive fields).

### Conclusion

Using a carefully designed psychophysical paradigm, combined with EEG and eye-tracking, our study demonstrates that induced alpha perceptual cycles travel across the retinotopic visual space in humans, at a propagation speed ranging from 0.3 to 0.5 m/s. These results suggest that brain activity travels across the visual cortex to modulate performance periodically across space and time.

## Data Availability Statement

Raw behavioral data and code relative to the main results will be available at https://github.com/CamilleFakche after publication.

## Author Contribution Statement

Conceptualization, C. F., L. D.; Formal Analysis, C. F.; Funding Acquisition, L. D.; Investigation, C. F.; Methodology, C. F., L. D.; Supervision, L. D.; Validation, L. D.; Visualization, C. F., L. D.; Writing – Original Draft Preparation, C. F.; Writing – Review and Editing, L. D.

## Acknowledgements

We thank Kirsten Petras, Andrea Alamia, Rufin VanRullen and Frédéric Chavane for their useful comments on the manuscript.

## Citation Diversity Statement

Retrospective analysis of the citations in every article published in this journal from 2010 to 2021 reveals a persistent pattern of gender imbalance: Although the proportions of authorship teams (categorized by estimated gender identification of first author/last author) publishing in the Journal of Cognitive Neuroscience (JoCN) during this period were M(an)/M = .407, W(oman)/M = .32, M/W = .115, and W/W = .159, the comparable proportions for the articles that these authorship teams cited were M/M = .549, W/M = .257, M/W = .109, and W/W = .085 (Postle and Fulvio, JoCN, 34:1, pp. 1–3). Consequently, JoCN encourages all authors to consider gender balance explicitly when selecting which articles to cite and gives them the opportunity to report their article’s gender citation balance. The authors of this article report its proportions of citations by gender category to be as follows: M/M = .642; W/M = .221; M/W = .074; W/W = .063.

